# Comparative analysis of anther development in natural populations indicate premature degradation of sporogenous tissue as a cause of male sterility in *Gaultheria fragrantissima*

**DOI:** 10.1101/368886

**Authors:** Wympher Langstang, Eros Kharshiing, Nagulan Venugopal

## Abstract

*Gaultheria fragrantissima* Wall. (Ericaceae) is a gynodioecious species having both hermaphrodite and male sterile plants. In this study, we present a comparative analysis of the different stages of anther development in naturally occuring hermaphrodite and male sterile populations of *G. fragrantissima* found in Meghalaya, India. While hermaphrodite flowers had well developed anther lobes, the male sterile flowers formed a white unorganized mass of tissues with a tuft of hairy outgrowth at the tip of the stamens. Histological analyses of progressive anther development in both the lines indicate an abnormal development of the sporogenous tissue in the developing anthers in the male steril line. While anther development in the hermaphrodite line was of the dicotyledonous type, the anthers of male sterile line showed progressive degradation of the sporogenous tissues and wall layers. Pollen development was also disrupted in male sterile line resulting in distorted pollen due to the irregular projection of exine wall. Our results suggest that premature degradation of the sporogenous tissues during anther development determines male sterility in *G. fragrantissima*.

## Introduction

In plants having diploid sets of chromosomes, microsporogenesis leads to the production of haploid spores by the meiotic division of a diploid microspore mother cell in the anther. During anther development, few hypodermal cells become differentiated into archesporial cells having larger size, slight radial elongation and conspicuous nuclei (Scott et al., 2004; Bhojwani 2015). These archesporial cells further divide to give rise to parietal cells towards the epidermis of the anther lobe and primiary sporogenous cells towards the interior of the anther. The primary sporogenous cells subsequently function as microspore mother cells resulting in pollen formation (Zhang and Yang 2014). In members of Ericaceae (including *Gaultheria*), microsporogenesis follows a similar mode of development except that the microspores are not individually separated and the resultant pollen are shed as tetrads (Davis 1966). The anther wall also varies from two to four cell layers, while the the tapetum is differentiated from an inner wall layer or from the sporogenous tissue (Batygina et al., 1963; Stushnoff and Palser, 1969; Herman and Palser 2000). The development of microscpore mother cells from the sporogenous is however concomitant with anther development (Venkateswarlu and Maheshwari 1973; Cambi and Herman 1989).

*G. fragrantissima* Wall. is an evergreen ericaceous shrub belongs to the family Ericaceae, distributed in South India, Indo–Malaya and North–East India. In North-East India, *G. fragrantissima* is found in the Shillong plateau at elevations of 1525 m above sea level (Haridasan and Rao 1985). *Gaultheria* which is a gynodioecious species has also been reported to exhibit male sterility in a majority of species. Male sterile plants have been reported in several species such as *G. colensoi* and *G. rupestris* var. subcorymbosa (Franklin 1962, Middleton 1991),; *G. crassca*; *G. rupestris* var rupestris (Delph 2006) and *G. fragrantissima* (Venugopal and Langstang 2011).

Male sterile mutants have been reported in a large number of plant species including that of the model plant *Arabidopsis* (Chaudhury, 1993; Dawson et al., 1993, Chase 2007). These include mutants defective in anther morphology, microsporogenesis, pollen development, and pollen function (Rick 1948; van Der Veen and Wirtz 1968; Albertsen and Phillips 1981; Kaul 1988; Regan and Moffat 1990). Many factors have been reported to contribute to male sterility in plants including variations in the abundance of amino acids in the locule (Fukasawa 1954; Brooks 1962), degeneration of the tapetum and alterations in tapetal morphology (Shi et al., 2009; Deng et al., 2018), premature degradation of callose wall (Worall et al., 1992; Wei et al., 2009) and meiotic defects during microsporogensis (Hooso et al., 2005). However, to our knowledge, the cause of male sterility and the developmental aspects of anther development in hermaphrodite and male sterile lines of *G. fragrantissima* have not been reported so far. Therefore, we investigated the developmental aspects of anther wall, sporogenous tissue, formation of pollen tetrads and ultrastructure of pollen grains in natural populations of hermaphrodite and male sterile plants of *G. fragrantissima* in an effort to determine the developmental basis of male sterility.

## Materials and Methods

### Plant material

For both hermaphrodite and male sterile lines used in this study, buds and opened flowers at different stages of development were periodically collected from naturally occuring populations found in Meghlaya, India (25°34^¢^0 N Latitude 91°52^¢^60 E Longitude, 1525 m asl). The collected buds and flowers were fixed in FAA [Formalin (5ml): Acetic acid (5ml): 70% Alcohol (90ml)], 3% Glutaraldehyde in 7.2 pH phosphate buffer and Cornoy’s fluid (Johansen 1940; Krishnamurthy 1988). The fixed samples were dehydrated using tertiary butyl alcohol series followed by impregnation with paraffin (Johansen 1940; Berlyn and Miksche 1976). The paraffin block were sectioned at 7-10 μm using Leica RM 2125 RT rotatory microtome. The resulting sections were deparafinised, rehydrated, stained and observed under a light microscope (Olympus BX43).

### Light microscopy studies

For light microscopy, the de-parafinised sections were rehydrated and stained with safranin and fast green, Sudan III (Johansen 1940). The sections were also stained with Periodic Acid Schiff’s reagent (Jensen 1962; Fedder and O’Brien 1968).

### Electron microscopy studies

For scanning electron microscopy the dissected anthers were fixed in 2-3% glutaraldehyde prepared in 0.1 M phosphate buffer (pH 7.2) at 4°C for 4 hrs. The samples were thoroughly washed in 0.1 M phosphate buffer and post fixed in 1% OsO_4_ for 2 hrs. After fixation, the plant material was dehydrated using increasing concentration of acetone. Dehydrated materials were dried in a Jeol JCPD-5 critical point dryer using 3-methyl butyl acetate solution as the exchange liquid. Dried materials were fixed on Eikon ion sputter, JFC-1100 and were coated with thin layer of gold vapour (300 Å layer). Gold coated plant materials were observed under Scanning electron microscope Joel JSM- 6360.

For transmission electron microscopy, samples were fixed in 2-3% glutaraldehyde prepared in 0.1 M phosphate buffer, pH 7.2 at 4°C for 8 hours, thoroughly washed in 0.1 M phosphate buffer and post fixed in 1% OsO_4_ for 2 hours. The samples were gradually dehydrated with acetone for 10-15 min in each step. Three changes were made in absolute acetone. The plant materials were embedded in Araldite CY 212. Ultrathin sections were cut at approximately 60-90 nm (600Å-900Å) through a Sorvall MT-2 ultramicrotome using diamond knife, and then stained with 2% aqueous uranyl acetate and lead citrate. Samples were observed under Zeiss EM-109 TEM.

## Results

### Sporogenous tissue is prematurely degraded in anthers of male sterile lines

During anther development in *G. fragrantissima* the tapetum persist until the formation of tetrads and remains intact until the maturation of the microspores (Chou 1952). In a previous report (Venugopal and Langstang 2011) it was observed that the flowers of the male sterile line of *G. fragrantissima* formed a white unorganized mass of tissues with a tuft of hairy outgrowth at the tip of the stamens, which was in stark contrast to that of the hermaphrodite line, which had well developed anther lobes having two distinct apical setaceous awns (Table 1). We then carried out histological analyses of progressive anther development in both hermaphrodite and male sterile lines in order to determine the developmental events that may be involved in the arrested anther development observed in the male sterile lines. In hermaphrodite flowers it was observed that the anther primordia were globular in outline which later appear rectangular due to accelerated cell division in four corners. In each corner of the anther primordium, 2-3 hypodermal cells become enlarged and differentiate into the archesporial initials, which are radially elongated containing dense cytoplasmic content and prominent large nucleus and nucleolus. Archesporial cells divide periclinally into an outer primary parietal cells and an inner primary sporogenous cells. The primary parietal cells further undergo periclinal divisions to form secondary parietal layers of which the inner secondary parietal layer divides and produces the middle layer and tapetum, whereas the outer one forms the endothecium. The primary sporogenous tissues are concomitantly formed along with anther wall development. The sporogenous tissues undergo several divisions and finally formed the microspore mother cells which are characterized by thin cell walls, dense cytoplasm and prominent nucleus. During the onset of meiosis in the microspore mother cell, a special callose wall layer is secreted around each microspore mother cell, which emits green yellowish fluorescence when observed under epifluorescent microscope. At the conclusion of meiotic divisions, the newly formed microscpores are arranged in tetrahedral and isobilateral condition resulting in the formation of tetrads (Figure 1a-c). Our observations thus indicate that anther development in the hermaphrodite line conforms to the dicotyledonous type as previously reported for members of Ericaceae including *G. fragantissima* (Cambi and Herman 1989; Hermann and Palser 2000). Anther differentiation in the male sterile line appears similar to that observed for the hermaphrodite line until formation of archesporial initials (Figure 1d). Here the sporogenous cells appear larger having dense cytoplasm and large nuclei. However, before the differentiation of wall layers, the sporogenous tissue divides abnormally to form central sterile septum. Subsequently the septum and the adjacent sporogenous tissue degraded totally which leads to complete degeneration of sporogenous tissues as well as wall layers (Figure 1e-g).

**Table 1:**
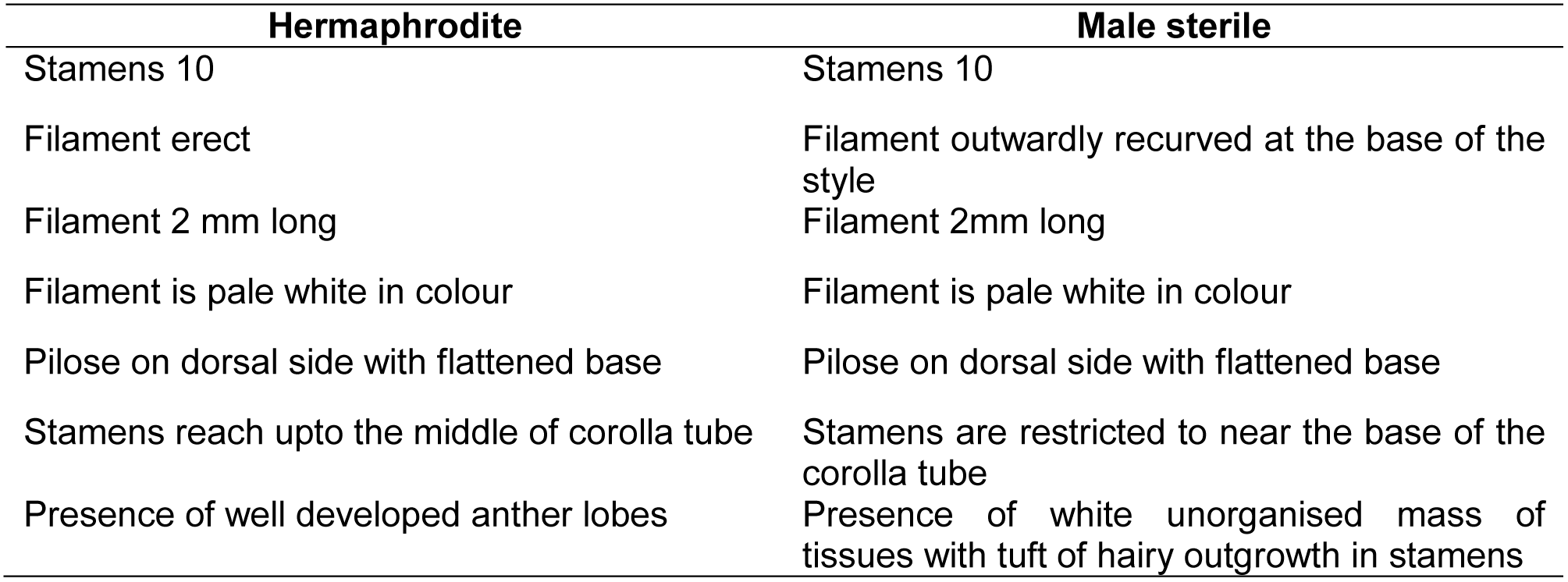
Comparison of stamen morphology between hermaphrodite and male sterile plants of *G.fragrantissima*

| **Hermaphrodite** | **Male sterile** |
| --- | --- |
| Stamens 10 | Stamens 10 |
| Filament erect | Filament outwardly recurved at the base of the style |
| Filament 2 mm long | Filament 2mm long |
| Filament is pale white in colour | Filament is pale white in colour |
| Pilose on dorsal side with flattened base | Pilose on dorsal side with flattened base |
| Stamens reach upto the middle of corolla tube | Stamens are restricted to near the base of the corolla tube |
| Presence of well developed anther lobes | Presence of white unorganised mass of tissues with tuft of hairy outgrowth in stamens |

**Figure 1:**
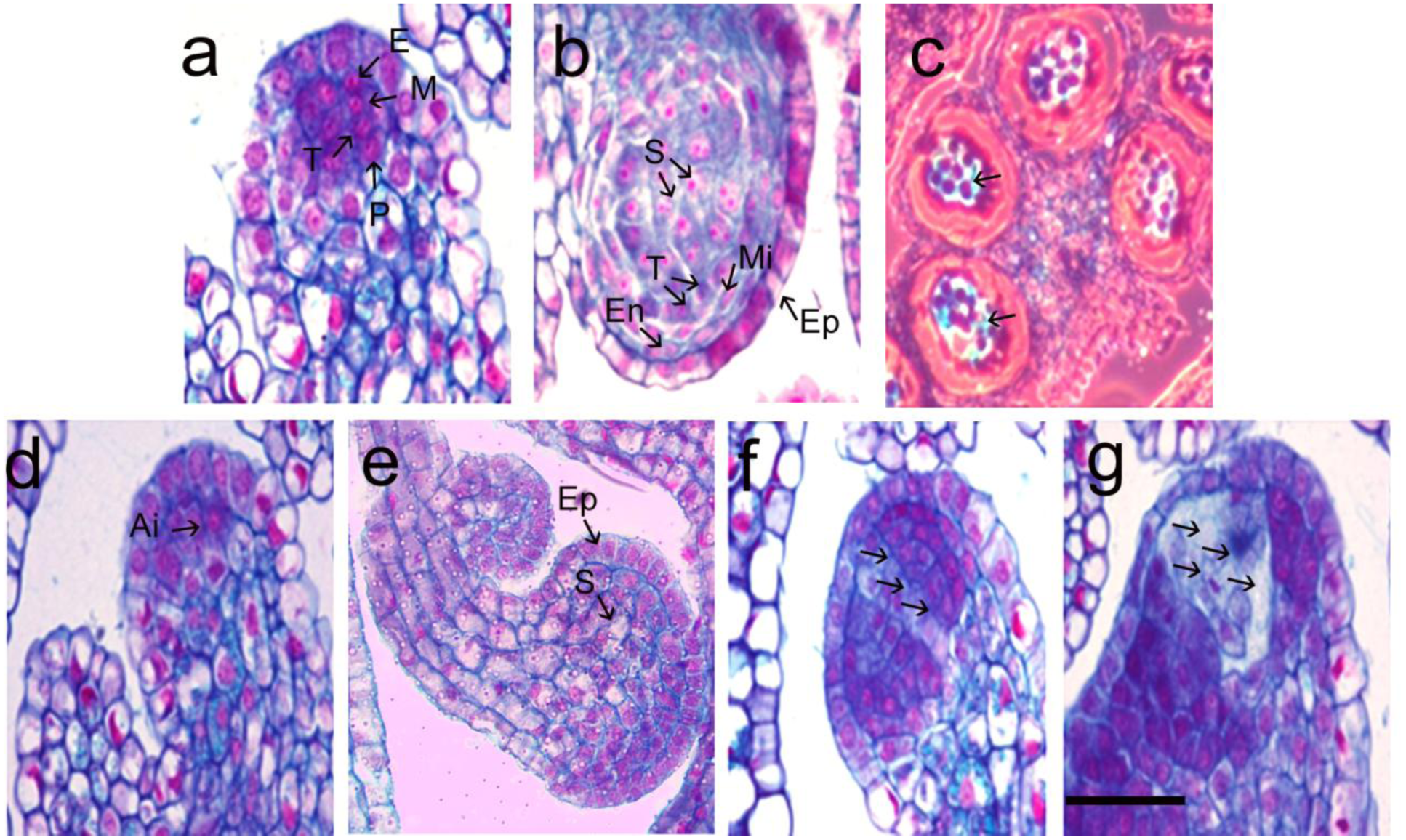
a,b,c: Microsporogenesis of hermaphrodite anther. a. Microsporangium of young anther lobe of hermaphrodite flower showing the initials for endothecium (E), middle layers (Ml), tapetum (T) and primary sporogenous initials. b. Microsporangium showing the tiered arrangement of epidermis (Ep),endothecium (En), middle layers (Ml) and tapetum (T) and central mass of sporogenous tissue (St). c. Transverse section of a single anther of hermaphrodite flower showing four lobes and the callose deposition around the microspore mother cell emit fluorescence under epifluorescence microscope stained with aniline blue (arrow). Scale bars – 260 μm. **d,e,f,g:** Microsporogenesis of male sterile anther.**d**. Microsporangium of young undifferentiated anther lobe of male sterile flower showing the normal differentiation of hypodermal archesporial initials (Ai). Scale bars – 23.58 μm.**e**. Microsporangium of a single anther of male sterile flower showing epidermis with abnormal development of archesporial initials. **f**. Showing the abnormal behavior of few archesporial initials leads to formation of central septum (S) within the anther lobe which divides the sporogenous tissue into halves. Scale bars – 20 μm.**g**. Showing the degradation of septum as well as anther wall layers and 99% degenerated sporogenous tissue. Scale bars – 20 μm.

### Male sterile line shows impaired pollen development

Transmission electron microscopic studies of pollen grains from hermaphrodite flowers indicate that the pollen are tricolpate with thick exine composed of sporopollenin and intine composed of pectic polymers with three distinct germinal pores. However, pollen of the male sterile flowers were distorted in shape due to the irregular projection of exine wall. The exine wall, which is primarily composed of sporopollenin, was also dissolved and disorganized possibly due to polymerization of sporopollenin. Furthermore while pollen from the male sterile lines showed distinct ektexine and endexine, similar to that in pollen of the hermaphrodite line, the the tectum and baculae of sexine in male sterile pollen were completely fused and developed a continous layer of radially oriented membranous granular material above the intine. Additionally, the outermost stratum of the intine was the most dense and compact and showed a defined microfibrillar structure (Figure 2, Table 2). Furthermore, while the central cytoplasm in normal pollen contained plastids with numerous starch grains, the central cytoplasm of pollen from male sterile line contained only 2-3 large starch grains.

**Figure 2:**
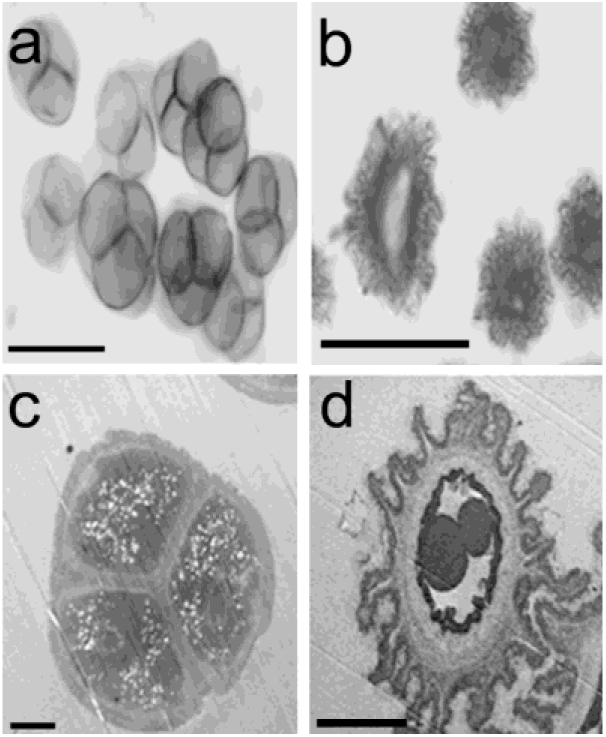
a: Emittence of blue fluorescence of tetrahedral pollen tetrad with aniline blue under epifluorescence microscope. Scale bars –12.5 μm. **b:** A few abnormal pollen grains differentiated with irregular projections of exine wall layer under light microscope. Scale bars – 20 μm. c,d: Transmission electron micrographs of mature pollen grains in microsporangia of hermaphrodite (c) anther showing showing the viable pollen tetrads of Hermaphrodite plant with distinct exine and intine. **Note-** abundance of cell organelles like proplastids, ribosomes, mitochondria, and electron translucent lipid droplets and electron dense bodies probably starch, protein etc. Scale bars 5 – μm. Scale bars –10 μm and male sterile (d) anther showing showing electron dense materials deposited below the intine (TEM); **Note-** the presence of starch grains and the extension of exine throughout the surface of the sterile pollen grain. Note the unusual enlargement of intine as well as thick cell wall. Scale bars – 2 μm.

**Table 2:** Comparison of pollen grain morphology between hermaphrodite and male sterile plant of *G.fragrantissima*

| **Hermaphrodite** | **Male sterile** |
| --- | --- |
| Pollen normal tetrahedral | Pollen irregular and distorted in shape |
| Pollen wall with distinct exine and intine | Pollen wall with irregular projection of exine. |
| Pollen with distinct ectexine and endexine. | Pollen with distinct ectexine and endexine. |
| Pollen cytoplasm contain plastids with numerous starch grains. | Pollen cytoplasm contain no plastids with 2-3 starch grains. |

## Discussion

The results of morphological and cytological investigations of microsporogenesis in hermaphrodite and male sterile lines of *G. fragrantissima* revealed broad spectrum deviations from the normal hermaphrodite anther development. In various plants, the morphological changes that precede male sterility occur at different developmental stages and tissues, ultimately manifesting as disturbed anther morphology. In male sterile lines of *Zinnia elegans*, male sterility occurs as a result of degenerate stamens, which are villous without pollen (Yao-mei et al., 2008), while in *Chrysanthemum morifolium* the anthers appear shrunken, brown-coloured and indehiscent (Li et al., 2010). In this study, we found that male sterile lines of *G. fragrantissima*, formed a white unorganized mass of tissues with a tuft of hairy outgrowth at the tips of the stamens resulting in unviable pollen. The anatomical characteristics of anther development in these male sterile lines is identical to that of the hermaphrodite anthers until the formation of archesporial initials. Anatomical studies relevant to male sterility, have reported several causes of male sterility such as tapetal abnormality (Holford et al., 1991, Strittmatter 2006, Chen et al., 2009), premature dissolution of callose (Worall et al 1992), anther (Dawson et al., 1993; Sanders et al., 1999) and meiotic defects (Arora and Gupta 1984, Graybosch and Palmer 1988; Zhang and Pan 1991). However, unlike previous reports, it is interesting to note that male sterility in *G. fragrantissima* may have occurred due to abnormal development of archespororial initials. Instead of cutting off parietal cells and primary sporogenous cells to form the wall layer and microspore mother cells respectively, the archesporial cells show precocious developmental arrest followed by collapse of the sporogenous tissues. In soybean, Stelly and Palmer (1982) reported arrest of anther development and subsequent degeneration of sporogenous tissues at all stages of anther development, from sporogenous cell stage to formation of pollen. Similarly in chives, cytoplasmic male sterility has been attributed to functional damages of sporophytic tissue (Engelke 2002). It therefore seems likely that similar to these reports, male sterility in *G. fragrantissima* reported in this study may have resulted from early degeneration of the sporogenous tissue.

During normal anther development, the archesporial initials cut off parietal cells toward epidermis to form the anther wall and primary sporogenous cells which contribute towards formation of microspores. The anatomical characteristics of anther development in male sterile lines of *G. fragrantissima* is identical to that of the hermaphrodite lines until the formation of archesporial initials. Thereafter the sporogenous tissue shows abnormal development resulting in premature degeneration. In some cases, it was observed that in few male sterile anthers, portions of the sporophytic tissue may have undergone division leading to formation of pollen. However, such pollen were observed to be abnormal and distorted. The pollen cytoplasm of such pollen was also condensed with disintegration of nuclear and vacuole membranes and most of the cell cytoplasm was occupied by starch grains. Similarly sterile pollen grains in *Helianthus annuus* are reported to have cytoplasm devoid of nucleus and possessing degenerated organelles with starch inclusions (Tripathi and Singh 2008). In *Citrus suavissima* mutants producing sterile pollens, Hu et al.,. (2007), also reported presence of thick electron dense material below the intine, similar to that observed in this study. We therefore propose that in *G. fragrantissima*, the abnormal development of the sporogenous tissues could be the likely cause of premature degeneration of sporogenous tissue and formation of non-viable pollen resulting in male sterility.

## Acknowledgement

This work was supported by Rajiv Gandhi National Fellowship, University Grants Commission, New Delhi to WL.

